# Structure of the malaria vaccine candidate Pfs48/45 and its recognition by transmission blocking antibodies

**DOI:** 10.1101/2022.05.24.493318

**Authors:** Kuang-Ting Ko, Frank Lennartz, David Mekhaiel, Bora Guloglu, Arianna Marini, Danielle J. Deuker, Carole A. Long, Matthijs M. Jore, Kazutoyo Miura, Sumi Biswas, Matthew K. Higgins

## Abstract

An effective malaria vaccine remains a global health priority and vaccine immunogens which prevent transmission of the parasite will have important roles in multi-component vaccines. One of the most promising candidates for inclusion in a transmission-blocking malaria vaccine is the gamete surface protein Pfs48/45, which is essential for development of the parasite in the mosquito midgut. Indeed, antibodies which bind Pfs48/45 can prevent transmission if ingested with the parasite as part of the mosquito bloodmeal. Here we present the first structure of full-length Pfs48/45, revealing its three domains to form a dynamic, planar, triangular arrangement. From this, we show where transmission-blocking and non-blocking antibodies bind on Pfs48/45. Finally, we demonstrate that antibodies which bind across this molecule can be transmission-blocking. These studies will guide the development of future Pfs48/45-based vaccine immunogens.

## Introduction

The recent approval of RTS,S as the first vaccine to prevent malaria caused by *Plasmodium falciparum* is an important step in efforts to end the devastating effects of this ancient disease [1], with promising results also recently released for the related R21 vaccine [2]. However, with large-scale trials of RTS,S indicating that it is around 30% effective at preventing severe disease [3], the quest for a more effective vaccine continues. The *Plasmodium* life cycle is complex and there are multiple stages at which such a vaccine could act [4], including fusion of the male and female gametes of the parasite within the mosquito midgut. If the bloodmeal of a mosquito contains antibodies which prevent gamete fusion, then these can stop completion of the parasite life cycle and can block transmission. Vaccine immunogens which induce the production of such transmission-blocking antibodies are therefore important components of multi-stage-targeting malaria vaccines [5].

One of the most promising transmission-blocking vaccine candidates is the gamete surface protein, Pfs48/45. This is essential for gamete fusion, as male *Plasmodium berghei* gametes that lack the homologous protein, Pbs48/45, are unable to penetrate female gametes to form zygotes [6]. In addition, antibodies induced by immunising animals with Pfs48/45 block the sexual development of the parasite within infected mosquitos [7-12]. Pfs48/45 is expressed on gametocytes found in human blood and is therefore exposed to the human immune system. Indeed, antibodies that target Pfs48/45 are also found in sera from individuals from malaria-endemic regions and the presence of such antibodies correlates with the transmission-blocking activity of these sera [13-17]. This means that individuals immunized with Pfs48/45 could also experience immune boosting through natural low-level infection. Also encouraging is the low sequence diversity of Pfs48/45 across strains of *Plasmodium falciparum* [12, 16, 18-20]. These factors combine to suggest that a vaccine immunogen based on Pfs48/45 will target a conserved and essential feature of the parasite life cycle and prevent further transmission from a malaria-infected individual.

As full-length Pfs48/45 has proved challenging to produce, vaccine immunogen design has so far been guided by an understanding of the molecular architecture of Pfs48/45 and by the identification of epitopes for potent monoclonal antibodies [5, 21]. Pfs48/45 is formed from three domains, with N- and C-terminal 6-cys domains joined by a central 4-cys domain, all linked to the gamete membrane through a C-terminal GPI anchor [22-24]. While these domain boundaries are well understood, how they come together to form full-length Pfs48/45 is still not known. Monoclonal antibodies have been generated against Pfs48/45, have been classified into different competition groups, and have been assessed for their transmission-blocking activity [8, 12, 25-28]. The most potent of these antibodies, 32F3 [8] and 85RF45.1 [25] bind to the C-terminal domain and the structure of the C-terminal domain bound to the Fab fragment of 85RF45.1 has been determined [12, 29]. This domain has therefore been the focus of most vaccine immunogen design approaches targeting Pfs48/45, primarily included as part of fusion proteins, such as that with GLURP [21]. However, other transmission-blocking antibodies have been identified which target the N-terminal and central domains of Pfs48/45 [12, 25, 28]. In this paper, we therefore present the structure of the full Pfs48/45 ectodomain, show how antibodies recognise different regions of Pfs48/45 and demonstrated that a significant fraction of the transmission-blocking activity of sera targeting Pfs48/45 is due to antibodies which target its N-terminal and central domains.

## Results

### A structural comparison of two antibodies targeting domain 3 of Pfs48/45

The two antibodies which have the highest transmission blocking activity, 32F3 and 85RF45.1, both bind to the C-terminal domain of Pfs48/45 [8, 12]. However, these two antibodies show substantial differences in transmission blocking activity, with 85RF45.1 completely preventing transmission at 14μg/ml, while 32F3 is only ∼40% effective at 70μg/ml [12]. To rationalise these differences, and to guide future vaccine design, we determined the structure of the C-terminal 6-cys domain of Pfs48/45 bound to 32F3 (Fig. 1A, Supplementary Table 1). We prepared Fab fragments from 32F3, combined with the Pfs48/45 C-terminal domain and conducted crystallisation trials. Crystals formed and a complete data set was collected to 1.9 Å resolution, allowing structure determination by molecular replacement.

**Figure 1:**
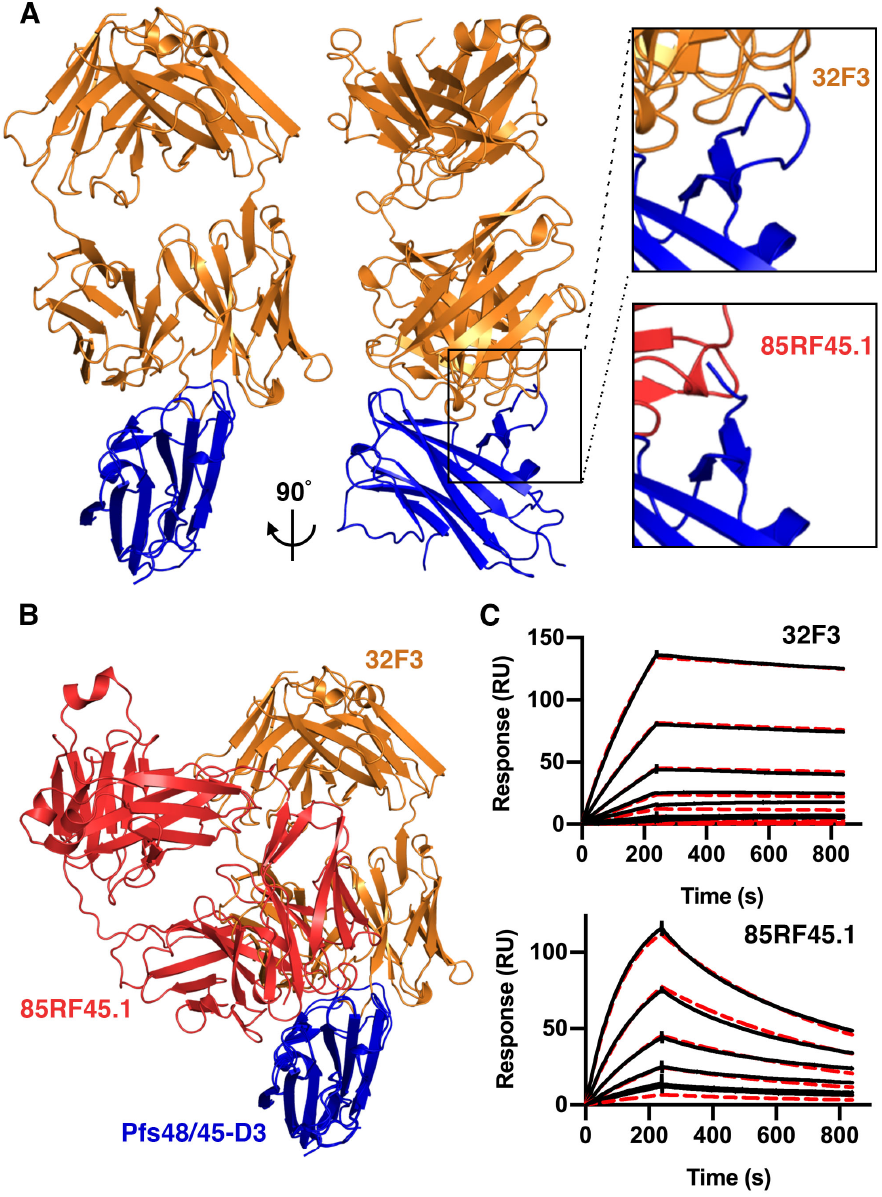
Comparison of the epitopes of transmission-blocking antibodies 85RF45.1 and 32F3. **A**. The structure of the C-terminal domain of Pfs48/45 (Pfs48/45-D3; blue) bound to antibody 32F3 (orange). The upper right inset shows a close-up of a Pfs48/45 loop which becomes ordered on 32F3 binding. The lower inset shows the equivalent view of the complex of Pfs48/45 (blue) bound to 85RF45.1 (red). **B**. An alignment of the structures of 32F3 (orange) and 85RF45.1 (red) bound to Pfs48/45-D3 (blue). **C**. Surface plasmon resonance analysis of the binding of Pfs48/45 binding to immobilised 32F3 and 85RF45.1. In both cases, the black lines show the responses due to a two-fold dilution series with a top concentration of 7.8nM for 85RF45.1 and 125nM for 32F3. The red dashed lines show fitting to a 1-to-1 binding model.

The structure of the C-terminal domain of Pfs48/45 was largely unchanged in conformation from that observed in complex with 85RF45.1, with a root mean square deviation of 0.66Å (Fig. 1A, B) [12, 29]. The most substantial change was in the loop comprising residues 357-369. This loop is disordered when bound to antibody 85RF45.1 but becomes ordered through interactions with 32F3 (Fig. 1A). While antibodies 32F3 and 85RF45.1 bind to overlapping epitopes, many of the contacts are not shared and the Fab fragments approach Pfs48/45 at different angles.

We also determined the binding kinetics for both 85RF45.1 and 32F3 to Pfs48/45, using surface plasmon resonance measurements (Fig. 1C, Supplementary Fig. 1). We first captured 85RF45.1 and 32F3 on different flow paths of a chip coated with protein A/G and flowed two-fold dilution series of Pfs48/45 over these antibody-coupled surfaces. Both antibodies bound to Pfs48/45 with similar affinities in the low nanomolar range (2.5nM for 85RF45.1 and 5.2nM for 32F3). However, binding kinetics differed, with 85RF45.1 showing faster association- and dissociation-rates, while 32F3 binds more slowly but forms a more stable complex. It seems likely that the slower binding kinetics of 32F3 may be due to the need for loop 357-369 to become ordered on binding, while antibody 85RF45.1 has an epitope which is unchanged in structure on antibody binding, allowing faster binding kinetics.

### Determining the structure of full-length Pfs48/45 by combining crystallography with an AlphaFold2 model

We next aimed to determine the structure of the complete Pfs48/45 molecule, allowing us to reveal the architecture of this three-domain protein and the locations of antibody epitopes outside the C-terminal domain. Insect cell expression systems were available to produce full-length Pfs48/45 ectodomain, or to generate a protein containing the central and C-terminal domains, Pfs4845-D2+3 [11]. Both were expressed without the GPI-anchor modification site and were purified and mixed in different combinations with one or more Fab fragments, selected from a set of antibodies which bind to different domains of Pfs48/45 (85RF45.1, 85RF45.3, 85RF45.5 [25], 32F3 [8], 10D8, 9D1, 7A6 [12]). Three of these complexes generated crystals: full-length Pfs48/45 bound to 10D8; full-length Pfs48/45 bound to both 10D8 and 85RF45.1; and Pfs4845-D2+3 bound to both 10D8 and 32F3. Data sets were collected to 4.2Å, 3.72Å and 3.69Å respectively.

We attempted to determine the structures of these complexes by molecular replacement, using the structures of the C-terminal domain bound to either 32F3 or 85RF45.1 Fab fragments as search models. In each case, the resultant electron density maps were sufficiently detailed to allow us to build a model for the central domain of Pfs48/45. This domain contains the epitope for 10D8, also allowing us to build a model for the 10D8 Fab fragment (Fig. 2, Supplementary Tables 1 and 2). However, while electron density could be observed for the N-terminal domain of Pfs48/45, this region was not sufficiently well resolved to allow a model to be built.

**Figure 2:**
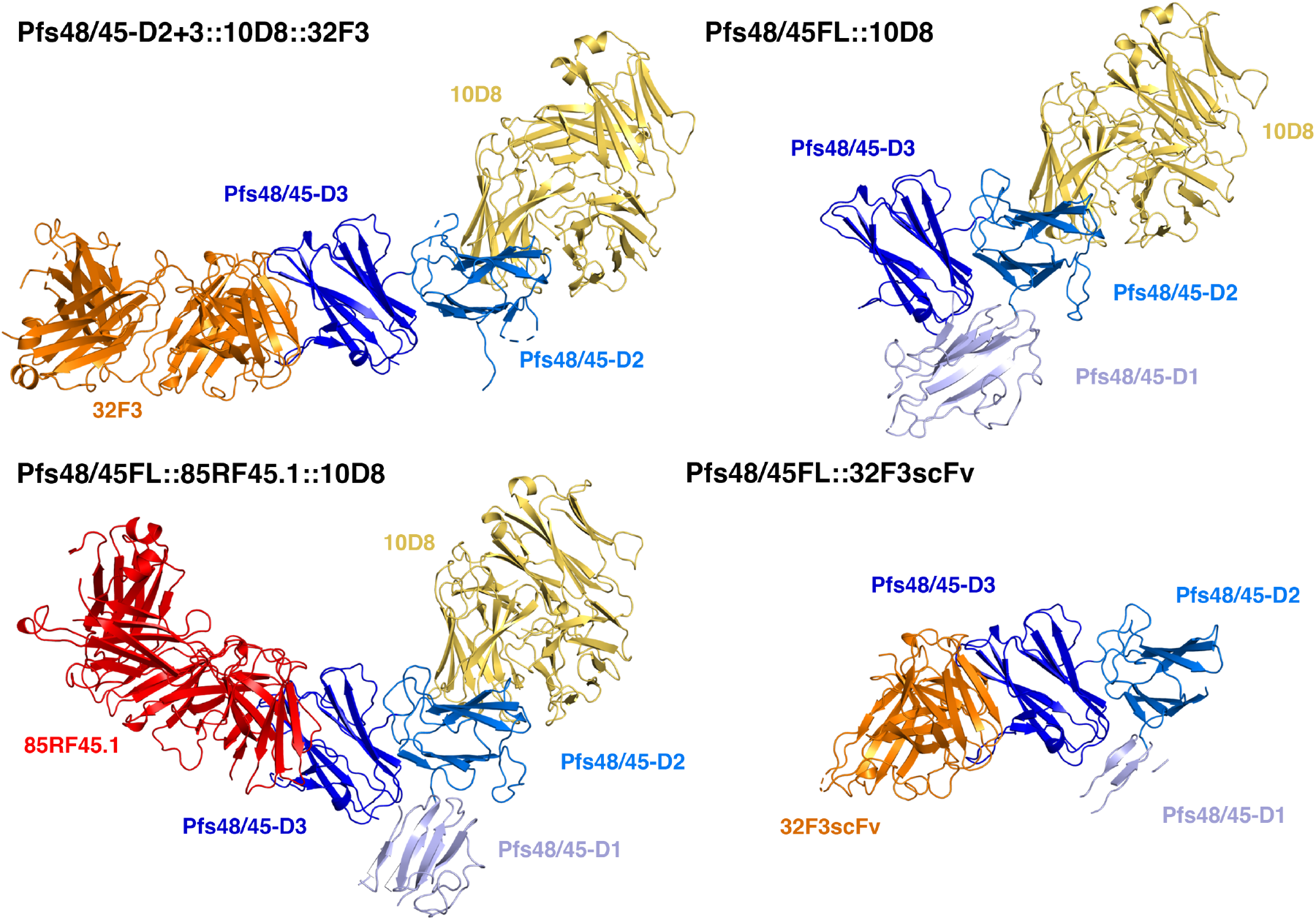
Structures of Pfs48/45 bound to antibodies 85RF45.1, 32F3 and 10D8. Structures of Pfs48/45 bound to different combinations of Fab fragments or scFvs. Pfs48/45 is shown with the three domains in different shades of blue. The N-terminal domain is light blue, the central domain is mid-blue and the C-terminal domain is dark blue. 10D8 is yellow, 85RF45.1 is red and 32F3 and its scFv fragment are orange.

The hinge between the variable and constant domains of Fab fragments is flexible. In case this flexibility reduced the degree of order within the crystals, we also generated scFv fragments for 85RF45.1, 32F3 and 10D8. These were purified, mixed with full-length Pfs48/45 and complexes were subjected to crystallisation trials. Crystals formed for the complex containing 32F3 scFv and a dataset was collected to 2.13Å resolution, allowing structure determination by molecular replacement. This higher resolution data allowed us to build an improved model for the central domain of Pfs48/45 (Fig. 2). However, the density for the N-terminal domain was still too poorly resolved to allow building of a complete molecular model of this domain. Analysis of packing within the different crystals suggested that, in each case, the N-terminal domain lies within a substantial cavity within the crystal lattice, and the lack of packing against other regions of Pfs48/45 allowed disorder of this domain, even within a well-ordered lattice.

Recent improvements in protein structure prediction from AlphaFold2 [30] provided a solution, allowing us to interpret the electron density maps for the N-terminal domain. AlphaFold2 correctly predicted the architecture of the central and C-terminal domains (with root mean square deviation of 1.26Å for the central domain and 0.43Å for the core 753/1044 atoms of the C-terminal domain), albeit with differences in loop structure (accession code Q8I6T1 at alphafold.ebi.ac.uk) (Supplementary Fig. 2). We therefore compared the AlphaFold2 model for the N-terminal domain with the electron density for this domain within the three different maps. The most complete electron density for this region of Pfs48/45 was obtained from crystals of full-length Pfs48/45 bound to 10D8. The AlphaFold2 model was therefore docked as single rigid body into this electron density and was rebuilt to fit the density. This generated a model for 122 residues of the 150 residue-long N-terminal domain, with a root mean square deviation of 1.63Å from the AlphaFold2 model. The N-terminus and loops 62-68 and 163-168 were unresolved. This model was then used to guide building of the observed portions of the N-terminal domain in full-length Pfs48/45 bound to 10D8 and 85RF45.1 (95 residues were built) and Pfs4845-D2+3 bound to both 10D8 and 32F3 (20 residues were built). Through this approach we generated the first molecular models of full-length Pfs48/45 derived from experimental crystallographic data, with an AlphaFold2 model used to guide building of the N-terminal domain (Fig. 2, Supplementary Tables 1-3).

### Pfs48/45 forms a dynamic three-domain disc-shaped architecture

We next used our structures, together with molecular dynamics simulations, to understand the organisation of Pfs48/45. Our three structures of Pfs48/45 all reveal a flattened, disc-like architecture (Fig. 3A). The C-terminal GPI-anchor membrane-attachment site is located on a flat surface of the molecule, suggesting that the disc may lie, on average, parallel to the membrane, with all three domains equally exposed at the gametocyte surface.

**Figure 3:**
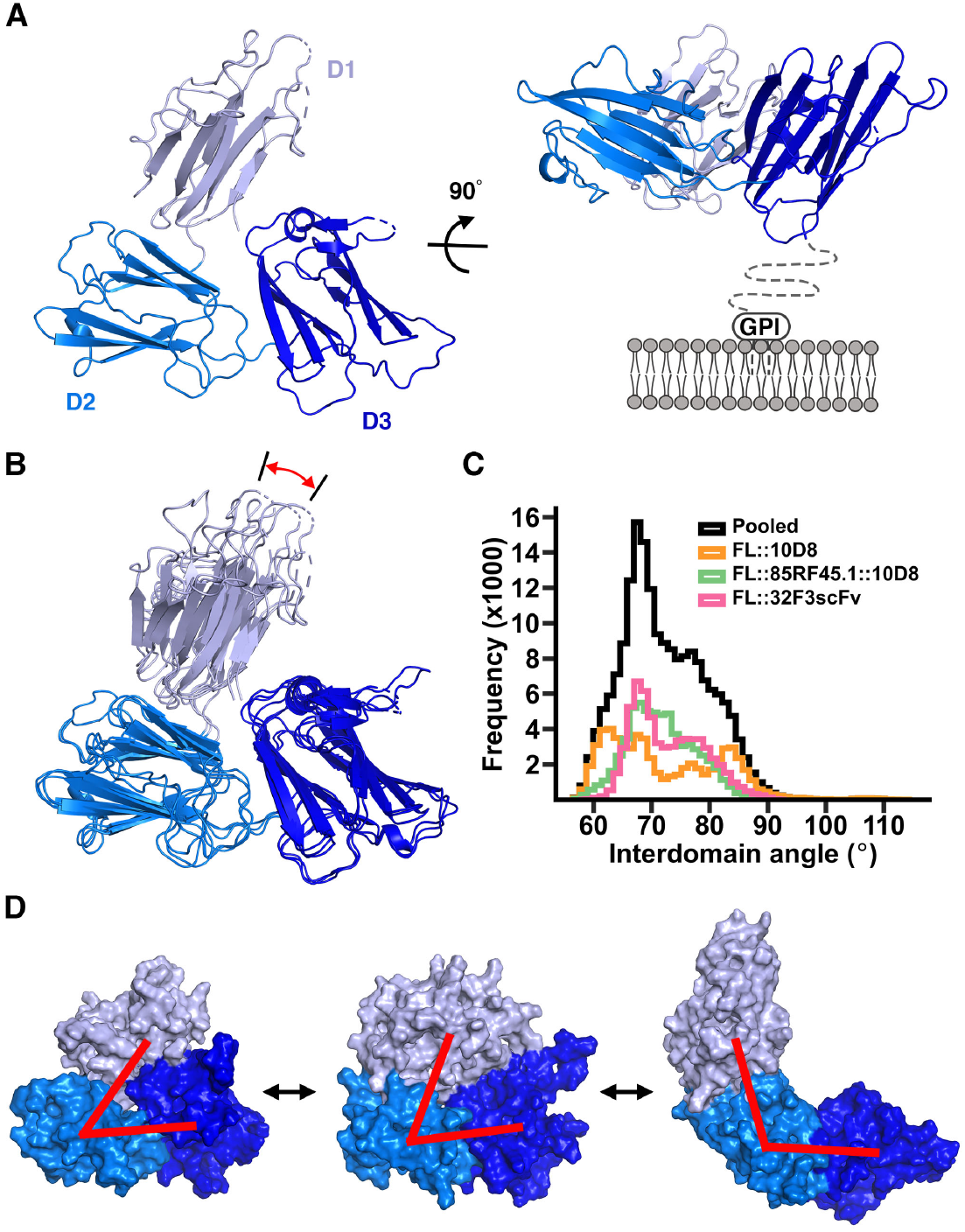
Structure and dynamics of Pfs48/45. **A**. The structure of Pfs48/45 taken from the structure of full-length Pfs48/45 bound to 10D8, viewed from two different directions. The N-terminal (D1), central (D2) and C-terminal domains (D3) are light, medium and dark blue. The right-hand panel also indicates the membrane and the 20 residue linker not included in our constructs. **B**. Comparison of the three full-length Pfs48/45 structures. Three structures were constructed by taking the models of Pfs48/45 from the three different crystal structures of Pfs48/45 bound to different antibody combinations and aligning the structure of the N-terminal domain onto the fragments of the domain built into the electron density. These composite structures have been aligned based on the central and C-terminal domains, showing the motion of the N-terminal domain, highlighted by the red arrow. **C**. Histogram showing the observed interdomain angles in full-length Pfs48/45 during atomistic molecular dynamics simulations. To define these angles two lines were drawn, linking the centres of mass of the N-terminal and central domains and those of the central and C-terminal domains. The angle shown is that between these lines. The three coloured histograms refer to outcomes of three independent simulations which started with the models in panel B with histograms being constructed by pooling five replicates for each starting structure. The black histogram is the pooled distribution of all replicates in all three starting models. **D**. The average angle observed in the simulations from C. is shown in the central panel with the most closed observed model in the left-hand panel and the most open model in the right hand panel. Red lines are as described in C.

The availability of three structures of Pfs48/45 allowed us to assess its degree of flexibility within crystals. We docked the most complete model of the N-terminal domain, from the structure of full-length Pfs48/45 bound to 10D8, onto the fragments of the domain observed in the two other crystal forms and the three resultant models for the full Pfs48/45 ectodomain were aligned (Fig. 3B). While the relative positions of the central and C-terminal domains were largely unchanged across these molecules, the position of the N-terminal domain changed, due to flexibility in the linker joining the N-terminal and central domains. Through these movements, the separation between the N- and C-terminal domains varied, with the vector of motion approximately parallel to the membrane axis.

To further assess the degree of motion within Pfs48/45 we used molecular dynamics simulations. Three separate simulations were run, with our three independent structures of Pfs48/45 containing the aligned N-terminal domain used as three distinct starting points. In each case, we simulated the system for 500 ns with 5 repeats. Analysis of these simulations revealed motion of the N-terminal domain in excess of that seen in the three crystal structures, leading to further separation of the N- and C-terminal domains. We plotted two lines though the structure; one linking the centres of mass of the N-terminal and central domains and one linking the centres of mass of the central and C-terminal domains. The angle between these lines was used as a measure of the degree of opening of the complex and varied from 56.4° to 114.9°, with an average of 72.8° (Fig. 3C, D, Supplementary Fig. 3). This compares with angles of 60.3°, 63.1° and 67.5° in the three crystal structures. Therefore Pfs48/45 is able to undergo substantial motion, through movement of the N-terminal domain, relative to the central and C-terminal domains, in a plane approximately parallel to the membrane. This flexibility may allow Pfs48/45 to adopt different conformations on binding to partners, such as Pfs230 and will also leave all three domains exposed to antibody recognition.

### Mapping transmission-reducing antibody epitopes to each domain of Pfs48/45

We next combined crystallography, electron microscopy and molecular dynamics simulations to assess how different monoclonal antibodies bind to Pfs48/45. In addition, to antibodies 32F3 and 85RF45.1 (Fig. 1), our crystal structures revealed the epitope for 10D8, an antibody with weak transmission-reducing potential which binds to the central domain of Pfs48/45 [12]. We combined the structure of Pfs48/45 bound to 10D8 with those of the C-terminal domain bound to 32F3 and 85RF45.1 to generate a composite model, showing the structure of Pfs48/45 bound to these three antibodies (Fig. 4A).

**Figure 4:**
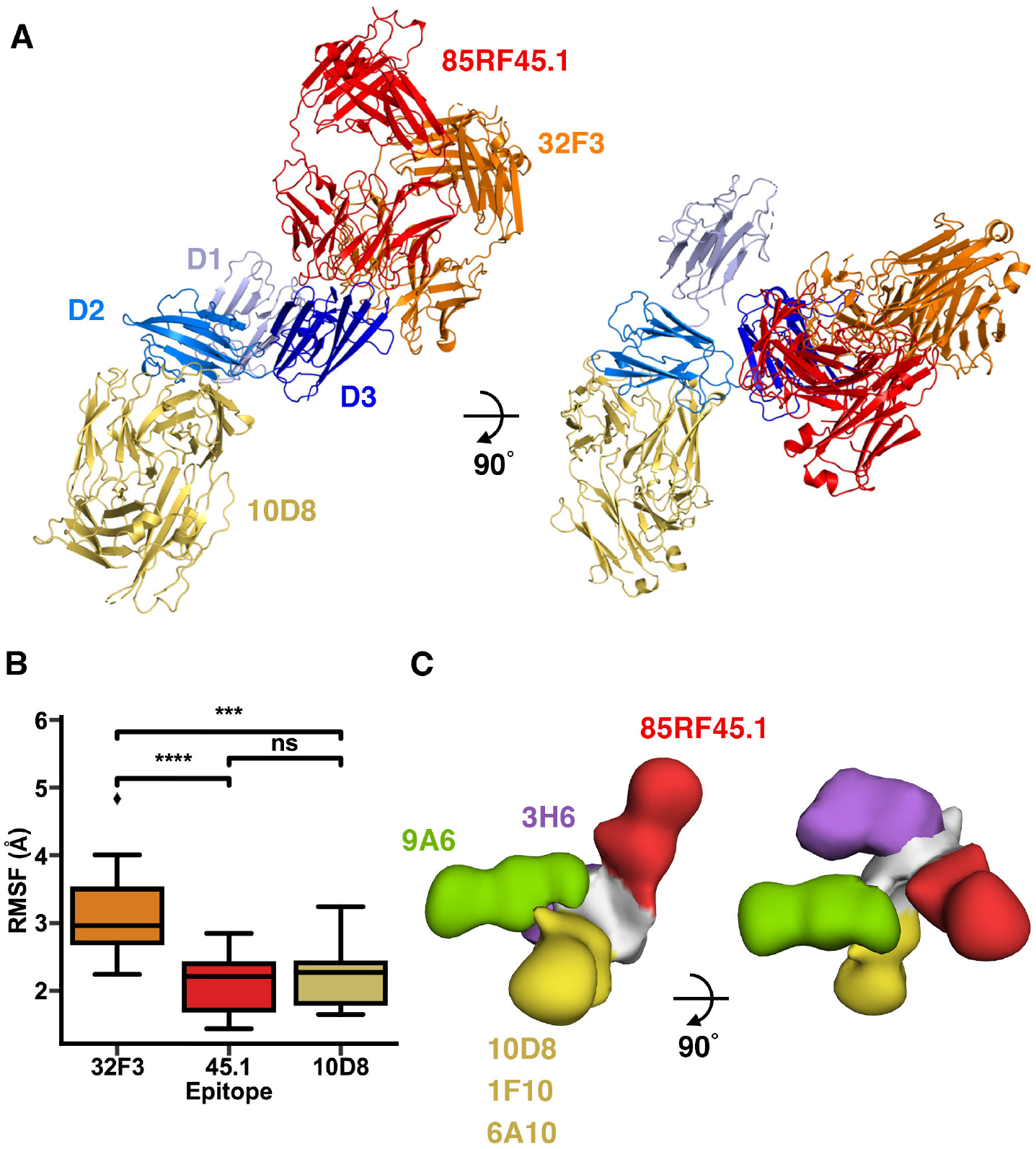
distribution of epitopes for transmission-blocking antibodies across Pfs48/45. **A**. The structure of Pfs48/45 bound to 10D8 is used as a template, and combined with structures of the D3 domain of Pfs48/45 bound to either 85RF45.1 or 32F3 to generate a model, showing the relative locations of the three antibody epitopes. **B**. A measure of the flexibility of the residues which form the epitopes for 32F3, 85RF45.1 and 10D8, as derived from molecular dynamics simulations. Each epitope was analysed in all 15 replicates and significance levels are based on Mann-Whitney U tests where *** indicates p<0.0005 and **** indicates p<0.00005. **C**. A composite model, derived from negative-stain electron microscopy structures. In each case, the complexes contained Pfs48/45 bound to 85RF45.1, together with either 9A6, 3H6, 10D8, 1F10 and 6A10. These were imaged by negative stain electron microscopy, and models were fitting into the resultant envelopes. These models were used to align the envelopes, allowing us to derive a composite model, showing the location for the five epitopes.

To understand the degree of flexibility of these three epitopes we analysed the molecular dynamics simulations presented in Figure 3, specifically assessing the motion of residues directly in contact with each of the three antibodies. This revealed that the epitope for 32F3 is significantly more flexible that those for 85RF45.1 and 10D8 (Fig. 4B), predominantly due to motion of the 357-369 loop, which forms part of the 32F3 epitope (Supplementary Fig. 3). This indicates that the slower association-rate for 32F3 compared to association rates for either 85RF45.1 or 10D8 is due to this flexibility and to the need for the epitope to adopt the correct structure on antibody binding.

Finally, we used negative stain electron microscopy to visualise the epitopes of four more antibodies. 1F10 and 6A10 are in the same competition group on the central domain as 10D8 while 9A6 is in a different competition group, binding to the same domain [12]. In contrast, 3H6 binds to the N-terminal domain. While 1F10 and 6A10 show some transmission-reducing activity, 9A6 and 3H6 do not [12]. In each case, we assembled complexes containing Pfs48/45, 85RF45.1 Fab and one other antibody Fab fragment, with 85RF45.1 included to provide a clear marker which could be used to align and position the additional antibody. We then used negative stain electron microscopy to image the complexes and single particle analysis to determine a low-resolution structure. The Pfs48/45:85RF45.1 complex and an additional Fab fragment model were then docked into these structures, taking into account to which domain the additional Fab bound to determine the organisation of the complex and the location of the Fab fragments. These five models were then aligned on Pfs48/45 and assembled together to show the locations of the five antibodies (Fig. 4C, Supplementary Fig. 4). The locations of 10D8, 1F10 and 6A10, which are part of the same competition group, were superimposable. In contrast, 9A6 and 3H6 adopted different locations on Pfs48/45, with both protruding in approximately the same plane as that shared by the three domains of Pfs48/45.

These findings suggest that both accessibility and also unknown functional properties of Pfs48/45 contribute to the transmission-blocking efficacy of antibodies which target Pfs48/45. When attached to the membrane, through a C-terminal GPI anchor, we would expect the Pfs48/45 disc to be on average arranged horizontal to the membrane plane (Fig. 3A). In this orientation, the epitope for 85RF45.1 will be most exposed on the membrane surface with the 32F3 epitope exposed to a lesser degree and the 10D8 epitope least accessible. This correlates with the order of transmission-blocking activity of these three antibodies, with 85RF45.1 as most potent and 10D8 as least potent. However, the epitopes for 9A6 and 3H6 emerge in the plane of Pfs48/45 and we would expect them to be more exposed on the membrane surface than the epitope for 10D8, and yet they are not transmission-blocking [12]. It is also notable that neither 3H6 or 9A6 stain gametocytes, suggesting these epitopes to be buried in that context [12]. A possible explanation for this is that the region of Pfs48/45 bound by these antibodies may be occluded by its binding partners, such as Pfs230. Further studies of Pfs48/45 function are required to determine whether this is the case.

### Antibodies targeting the N-terminal and central domains of Pfs48/45 substantially contribute to transmission-blocking activity of sera

The structure of Pfs48/45 indicates that each of its three domains will be exposed on the gametocyte surface (Fig. 3) and antibodies binding to each domain have been shown to stain gametocytes [12]. However, the most effective known transmission-blocking monoclonal antibodies bind to the C-terminal domain [8, 25]. We therefore aimed to determine the degree to which antibodies against the N-terminal and central domains contribute to transmission-blocking activity, using a combination of immunisation and antibody depletion experiments.

We first produced protein consisting of the N-terminal and central domains of Pfs48/45 (Pfs48/45-D1+2) and used this to immunise mice. IgG was purified from these mice and the titres of antibodies targeting Pfs48/45-D1+2 were determined by end-point ELISA (Fig. 5A). The transmission-blocking activity of these antibodies was then assessed using a standard membrane feeding assay. Antibodies were added to blood containing *Plasmodium falciparum* gametocytes, this was fed to mosquitos and the number of ookinetes formed in mosquito midguts was counted. Total IgG, purified from mice immunised with either 0.1μg or 1μg of Pfs48/45-D1+2 was tested at a concentration of 750μg/ml, causing almost complete inhibition (96% reduction) in oocyte numbers (Fig. 5B). When tested at 375 and 188μg/ml, the total IgG from the 1μg group also showed significant inhibition of 92% and 68% respectively (Fig. 5C). Therefore, protein immunogens containing just the N-terminal and central domains of Pfs48/45 are able to induce highly potent transmission-blocking antibodies.

**Figure 5:**
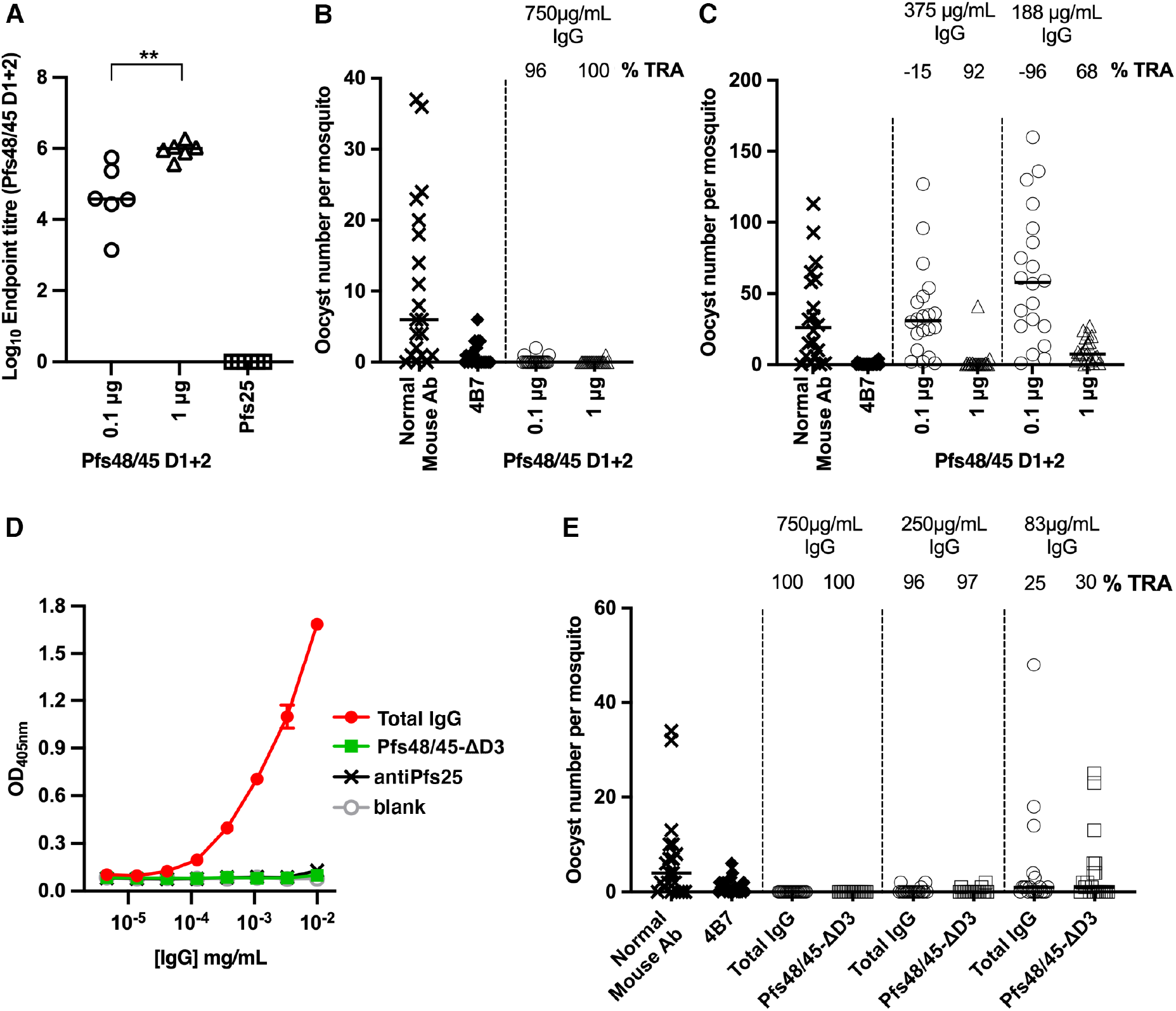
Transmission-blocking antibodies target the N-terminal and central domains of Pfs48/45. **A, B, and C**. Female CD1 mice were immunised twice with either 0.1µg or 1µg of Pfs48/45-D1+2, or 1µg of Pfs25. Sera was collected three weeks after the final dose for analysis. **A)** IgG titres as measured by endpoint ELISA using Pfs48/45-D1+2 as the coating antigen. Each symbol is for the serum sample from an individual mouse; lines represent the median of each group. Mann-Whitney 2-tailed test was performed to compare the two Pfs48/45-D1+2 groups (p = 0.0043). **B,C**. Transmission-blocking efficacy of IgG induced by immunisation with Pfs48/45-D1+2. Total IgG was purified from the pooled serum of each group (3 weeks post-boost) and mixed with *P. falciparum* NF54 cultured gametocytes at 750µg/mL (**B)** and in a separate experiment at 375µg/mL and 83µg/mL (**C**) and fed to *A. stephensi* mosquitoes (n = 20 per test group). IgG from naive mice was used as a negative control (“normal mouse Ab”); the transmission blocking anti-Pfs25 mAb 4B7 was used as a positive control. Data points represent the number of oocysts in individual mosquitoes 8 days post-feeding; lines show the arithmetic mean. **D** and **E**. Female CD1 mice were immunised three times with 5µg of Pfs48/45-FL. Three weeks after the final dose sera were collected and total IgG purified. Total IgG was depleted of Pfs48/45-D3-targeting IgG using a column coupled with Pfs48/45-D3, resulting in Pfs48/45-ΔD3. **D**. Pfs48/45-D3 specific ELISA showing lack of recognition of Pfs48/45-D3 by Pfs48/45-ΔD3. Pfs25 specific IgG (αPfs25) was included as a negative control. **E**. Transmission-blocking efficacy of total IgG from Pfs48/45-FL or Pfs48/45-ΔD3 fed to *A. stephensi* mosquitoes at concentrations of 750µg/mL, 250µg/mL and 83µg/mL (n = 20 per test group). IgG from naive mice was used as a negative control (“normal mouse Ab”); transmission blocking anti-Pfs25 mAb 4B7 was used as a positive control. Data points represent the number of oocysts in individual mosquitoes; lines show the arithmetic mean. (TRA, Transmission Reducing Activity [% inhibition in mean oocyst count per mosquito]).

We next used a depletion approach to assess the contribution that antibodies which target the N-terminal and central domains of Pfs48/45 make to the transmission-blocking activity of IgG from mice immunised with full-length Pfs48/45. Total IgG from mice immunised with 5μg of full-length Pfs48/45 was tested at a concentration of 750μg/ml and showed 100% transmission blocking activity (Fig. 5E). This IgG was depleted of antibodies which bind to the C-terminal domain of Pfs48/45 using an affinity column and was tested for transmission-blocking activity (Fig. 5D). At all three total IgG concentrations tested, the sera depleted of C-terminal domain-targeting antibodies showed no significant difference to the sera prior to depletion (Fig. 5E). This shows that the majority of the antibodies with transmission-blocking activity induced through immunisation with full-length Pfs48/45 bind to either the N-terminal or central domains.

## Conclusions

The structure of the Pfs48/45 ectodomain reveals that its three domains are arranged as a disc. The location of the C-terminal GPI-anchor attachment site on one of its flatter surfaces will cause the disc to lie approximately parallel to the membrane plane, with all three domains accessible on the cell surface. The presence of a flexible linker between the N-terminal and central domains also causes the disc to be dynamic, due to opening and closing of the gap between the N- and C-terminal domains. How this structure and dynamics relate to the function of Pfs48/45 is currently unclear. Pfs48/45 on gametocyte surfaces is part of a larger protein complex including other proteins essential for gamete fusion, including Pfs230 [31] and the PfCCp proteins [32, 33]. The molecular architecture of this complex and the mechanistic roles of its components during gamete fusion, are currently not known. The structure of Pfs48/45 makes it tempting to speculate that the open, dynamic membrane-distal surface of the disc, as presented on gametocytes, act as a platform that allows it to bind important interaction partners such as Pfs230 and the PfCCp proteins. To test this will require further study, both to determine how this complex is arranged and to understand which parts of Pfs48/45 are occluded by binding partners.

The structure of Pfs48/45 also has implications for our understanding of what makes an effective transmission-blocking antibody. A number of criteria might determine the potency of such an antibody, including the location of its epitope relative to functionally important sites on the protein, its binding kinetics and the accessibility of its epitope when presented on the parasite surface. Our lack of mechanistic insight into the role of Pfs48/45 during gamete fusion currently makes it impossible to determine whether the efficacy of antibodies correlates with their effects on protein function. However, our structural characterisation of antibody epitopes, coupled with kinetic data and measurements of transmission-blocking activity, suggest that epitope accessibility is a key feature of highly potent antibodies. In particular, comparison of 85RF45.1 and 32F3 reveals that these two antibodies have overlapping epitopes on the C-terminal domain and have very similar affinities. The flexible nature of the 32F3 epitope, compared with the more rigid epitope for 85RF45.1 affects their kinetics of binding to soluble Pfs48/45 protein, with 85RF45.1 binding and dissociating more rapidly. However, unlike antigens involved in erythrocyte invasion [34], Pfs48/45 is exposed for significant periods of time on the gamete surface, making it unlikely that fast association-rates are important for antibody efficacy. Instead, the most likely cause of differences in potency of 85RF45.1 and 32F3 is the degree to which their epitopes are exposed on the gametocyte surface. While 32F3 approaches from the side of the disc, 85RF45.1 binds on the more exposed, membrane-distal surface, allowing greater accessibility to its epitope in the context of the gametocyte surface. Indeed, transmission blocking antibody 10D8 has lower efficacy than either 85RF45.1 and 32F3 and binds to an epitope in a less exposed position on the Pfs48/45 surface. However, exposure and accessibility does not provide the full story. Antibodies 3H6 and 9A6 bind to more exposed surfaces of Pfs48/45 than 10D8 and yet are not transmission blocking and do not bind to gametocytes, most likely due to occlusion by Pfs48/45 binding partners. Future studies are needed to reveal how its interactions with other binding partners modulates its activity and affect vaccine design decisions.

Finally, our studies of full-length Pfs48/45 have consequences for vaccine design. Many current approaches for Pfs48/45 have focused on the C-terminal domain, which is easier to produce than full-length Pfs48/45 and contains the epitopes for the most potent known transmission-blocking antibodies. However, the structure of Pfs48/45 shows that all three of its constituent domains will be exposed when membrane anchored. Our depletion studies also show that the majority of transmission-blocking antibodies raised through immunisation of mice with full-length Pfs48/45 bind to the N-terminal and central domains. There are still many uncertainties about the role of Pfs48/45 during gamete fusion, including when Pfs48/45 assembles into a complex with other gametocyte surface proteins or which epitopes are exposed when it is part of this complex. However, our current data suggests a vaccine design strategy which includes full-length Pfs48/45, to take advantage of epitopes for transmission-blocking antibodies which target the N-terminal and central domains, perhaps first generating versions of Pfs48/45 which are thermally stabilised to prevent dynamic movement. It also indicates the importance of correct presentation of Pfs48/45 during vaccination. Immunisation with soluble, or incorrectly oriented Pfs48/45, will induce antibodies which target epitopes poorly accessible on the gametocyte surface. In contrast, a vaccine particle which presents Pfs48/45 in an orientation consistent with that on the gametocyte surface is more likely to induce antibodies, like 85RF45.1, with potent transmission-blocking potential.

### Experimental procedures

#### Pfs48/45 expression and purification

The sequences for all Pfs48/45 protein variants (PlasmoDB:PF3D7_1346700: Pfs48/45 full length residues 27-427, Pfs48/45-D1+2 residues 27–288, Pfs48/45-D2+3 residues 159-428 and Pfs48/45-D3 residues 291-428) were codon optimised for expression in Drosophila melanogaster (GeneArt Life Technologies), (GeneArt Life Technologies), with a Drosophila BiP signal peptide introduced at the N-terminus and the four amino acids EPEA (C-Tag) inserted at the C-terminus. All N-glycosylation sites were left intact. The sequence was subcloned into the Drosophila S2 expression vector pExpres2.2 (ExpreS2ion Biotechnologies). Polyclonal Drosophila S2 stable cell lines were generated by non-viral transfection into ExpreS2 Drosophila S2 cells (Expres2ions Biotechnologies) with selection via G418 resistance. Recombinant Pfs48/45-D1+2, Pfs48/45-D2+3 and Pfs48/45-D3 were purified by initially concentrating the cell culture supernatant using a Tangential Flow Filtration system with a Pellicon 3 Ultracel 3 kDa membrane (Merck Millipore, UK). Pfs48/45 full-length cell culture supernatant was concentrated as above but using a Pellicon 3 Ultracel 10 kDa membrane (Merck Millipore, UK). The concentrated supernatants were then loaded onto CaptureSelect™ C-tag affinity columns (Thermo Fisher Scientific) equilibrated in Tris-buffered saline and bound proteins were eluted with 20 mM Tris–HCl, 2 M MgCl2, pH 7.4. Protein containing fractions containing were then pooled and subjected to size-exclusion chromatography using a Superdex 200 16/600 PG column (Cytiva). Proteins were aliquoted and flash frozen in liquid nitrogen for later use.

#### Expression and purification of antibodies and scFvs

32F3, 85RF45.1, 10D8, 1F10, 3H6, 6A10 and 9A6 were expressed from hybridoma cell lines, purified using protein A/G and ficin cleaved to generate Fab fragments, as described [12]. To produce scFv fragments, the sequence of 32F3 scFv was designed with the variable region of the heavy chain (from residue DVKLV to residue TLTVS) followed by an 18-residue linker (GGSSRSSSSGGGGSGGGG) and the variable region of light chain (from residue QIVLS to KLELK). The sequence of the 85RF45.1 scFv contains the variable region of the light chain (from residue QFVLS to residue KLTVT), followed by the 18-residue linker and the variable region of the heavy chain (from residue EVQLV to residue MVTVS). The sequence of 10D8 scFv contains the variable region of the light chain (from residue DIVMS to residue TKLEI), followed by the 18-residue linker and the variable region of the heavy chain (from residue EVMLV to residue GTSVT). Codon optimised synthetic genes (ThermoFisher) of 10D8 scFv, 32F3 scFv, and 85RF45.1 scFv were PCR amplified and subcloned into a pHLsec vector containing a C-terminal hexa-histidine tag. These were expressed in HEK293 cells. Cultures were harvested six days after transfection. The supernatant was filtered and incubated with Excel Ni Sepharose resin (GE Healthcare) at 4°C for 30 minutes. The mixture was then applied onto a gravity flow column. The column was washed with 20mM Tris pH 8.0, 150mM NaCl, 10 mM imidazole and eluted in the same buffer containing 250 Mm imidazole. Elution fractions were collected and concentrated to run gel filtration with superdex 200 increase 10/300 (GE Healthcare) in 20mM Tris pH 8.0, 150mM NaCl.

#### Crystallisation and structure determination

Complexes containing different combinations of full-length Pfs48/45, the C-terminal domain of Pfs48/45 (Pfs48/45-D3) and the central and C-terminal domains (Pfs48/45-D2+3), with one or more Fab fragment or scFvs, were prepared by mixing different components in stoichiometric ratios. Complexes were purified by size exclusion chromatography on two Superdex Increase 200 10/300 columns (GE healthcare) in series into 10mM HEPES, 150mM NaCl, pH 7.2. Peak fractions containing pure complex were then concentrated and used to set up vapour diffusion crystallisation trials in sitting drops by mixing 100nl of protein complex with 100nl of well solutions.

Crystals formed in the following conditions. For Pfs48/45-D3:32F3, initial crystals formed at a protein concentration of 11.1 mg/ml at 293K in 0.04M potassium dihydrogen phosphate, 16% (w/v) polyethylene glycol (PEG) 8000 and 20 % (v/v) glycerol. Crystallisation conditions were further optimised by increasing the concentration of polyethylene glycol 8000 to 25% (w/v) and omitting potassium dihydrogen phosphate during crystallisation. For data collection, these crystals were transferred into well solution containing 30% (v/v) glycerol and flash-cooled in liquid nitrogen. For Pfs48/45-D2+3:10D8:32F3, crystals formed at a protein concentration of 10.5 mg/ml at 293K in 10 % (w/v) PEG monomethyl ether 5000, 0.1M 2-(N-morpholino) ethanesulfonic acid pH 6.5 and 12% (v/v) 1-propanol. Initial crystals were further optimised by microseeding into the same condition. For data collection, crystals were transferred into well solution containing 25% (v/v) 2-Methyl-2,4-pentanediol and flash-cooled in liquid nitrogen. For Pfs48/45-FL:10D8, the best crystals grew at a protein concentration of 10.7 mg/ml at 277K in 10 % (w/v) PEG 1000 and 10% (w/v) PEG 8000. For data collection, crystals were transferred into well solution containing 25% (v/v) 2-Methyl-2,4-pentanediol and flash-cooled in liquid nitrogen. Crystals for Pfs48/45-FL:85RF45.1:10D8 formed at a protein concentration of 13.4 mg/ml at 277K in 0.1M Lithium sulfate, 0.1M HEPES pH 7.0 and 20 % w/v Polyvinylpyrrolidone. For data collection, crystals were transferred into well solution containing 30% (v/v) glycerol and flash-cooled in liquid nitrogen. For Pfs48/45FL:32F3 scFv, crystals formed at protein concentration of 12 mg/ml at 291K in 0.1M MES pH6.0, 0.2M CaCl_2_, 20% (w/v) PEG6000 and for data collection crystals were transferred into well solution containing 25% (v/v) glycerol.

Data were collected on the following beamlines. Data for Pfs48/45-D3:32F3 and for Pfs48/45-FL:10D8 were collected at IO4 (Diamond Light Source, UK); data for Pfs48/45-D2+3:10D8:32F3 were collected at IO4-1 (Diamond Light Source, UK); data for Pfs48/45-FL:85RF45.1:10D8 were collected at IO3 (Diamond Light Source, UK) and data for Pfs48/45-FL:32F3 scFv were collected at IO4 (Diamond Light Source, UK). The data for Pfs48/45-FL:85RF45.1:10D8, Pfs48/45-D3:32F3, Pfs48/45-D2+3:10D8:32F3 and Pfs48/45-FL:10D8 were processed with the CCP4i2 programme suite [35] using the Xia2/DIALS pipeline [36]. For Pfs48/45FL:85RF45.1:10D8, two datasets from the same crystal were merged together. Data for Pfs48/45-FL:32F3 scFv were processed with XDS. All structures were determined by molecular replacement using Phaser-MR [37] with model building in Coot [38] and refinement using phenix.refine [39] and Buster [40]. All structure figures were prepared using PyMol (Schroedinger LLC).

#### Surface plasmon resonance analysis

Surface plasmon resonance experiments were carried out using a Biacore T200 instrument (GE Healthcare), to analyse binding of Pfs48/45 to 85RF45.1, 32F3 and 10D8. Pfs48/45 was buffer-exchanged into 20mM HEPES pH 7.2, 300mM NaCl, 0.05% Tween-20. Individual mAbs were immobilised on a CM5-chip (GE Healthcare) pre-coupled to Protein A/G (Thermo Fisher) and Pfs48/45 was injected over the chip surface at a flow rate of 30µl/min, with 240s association time and 600s dissociation time. For all three antibodies, we injected a series of samples, forming two-fold dilution series, with a top concentration of 7.8nM for 85RF45.1 and 125nM for 32F3 and 10D8. After each injection, the chip surface was regenerated with 10mM Glycine, pH 2.0 for 120s at 10µl/min, followed by a regeneration period of 180s. Data were analysed using the BIAevaluation software 2.0.3 (GE Healthcare).

#### Molecular Dynamics Simulations

To prepare structures for simulation, we first added models for the missing regions in the Pfs48/45-FL:85RF45.1∷10D8 and Pfs48/45-FL:32F3 scFv structures by grafting the model for the N-terminal domain from the Pfs48/45-FL:10D8 structure on the existing partial N-terminal domain models in the former two structures. Next, we modelled missing loops in each structure individually using MODELLER (v. 10.2) [41]. We allowed refinement only in the missing loops and generated 1000 decoys corresponding to residues 45-428 for each structure and picked the final models using the SOAP [42] scoring function. We used OpenMM (v. 7.7) [43] to run atomistic molecular dynamics simulations for each starting structure. To avoid non-physiological interactions due to truncation, we capped N- and C-termini using acetyl and amide groups, respectively. Next, we protonated the models at a pH of 7.5, soaked them in truncated octahedral water boxes with a padding distance of 1 nm, and added NaCl to an ionic strength of 150 mM to neutralise charges. We parameterised the systems using the Amber14-SB force field [44] and modelled water molecules using the TIP3P-FB model. Non-bonded interactions were calculated using the particle mesh Ewald [45] method using a cut-off of distance of 0.9 nm, with an error tolerance of 0.0005. Water molecules and heavy atom-hydrogen bonds were rigidified using the SETTLE [46] and SHAKE [47] algorithms, respectively. We used hydrogen mass repartitioning [48] to allow for 4 fs time steps. Simulations were run using the mixed-precision CUDA platform in OpenMM using the Middle Langevin Integrator and the Monte-Carlo Barostat. We equilibrated systems using a multi-step protocol: (i) energy minimisation over 10000 steps, (ii) heating of the NVT ensembles from 100 K to 300 K over 200 ps, (iii) 200 ps simulation of the NPT ensembles at 300 K, (iii) cooling of the NVT ensembles from 300 K to 100 K over 200 ps, (iv) energy minimisation over 10000 steps, (v) heating of the NVT ensembles from 100 K to 300 K over 200 ps, and (vi) 5 ns simulation of the NPT ensembles at 300 K. Following this, we ran production simulations of NPT ensembles for 500 ns. We used MDTraj (v. 1.9.6) [49] for analyses of resulting trajectories.

#### Negative stain cryo-electron microscopy and image processing

Each Fab was mixed with Pfs48/45-FL and 85RF45.1 Fab at the ratio of 1:1:1 and incubated on ice for 30 minutes in 20mM Tris pH 7.5, 150mM NaCl. The mixture was then injected onto Superdex 200 Increase 10/300 column (Cytiva) for size-exclusion chromatography. Fractions corresponding to Pfs48/45-FL bound to two Fabs (around 150 kD) were collected and concentrated. Purified complexes were flash-frozen using liquid nitrogen and stored at - 80°C.

For negative staining, purified complexes of Pfs48/45:85RF45.1 Fab with one additional Fab, from 1F10 Fab, 3H6 Fab, 6A10 Fab, 9A6 Fab or 10D8 Fab were diluted to 10 µg/ml. Carbon support 200-mesh copper grids were glow-discharged at 15 mA for 20 s. 5 µl protein sample was applied onto the grids for 60 s and blotted. Then the grids were washed twice with 20mM Tris pH 7.5, 150mM NaCl buffer and stained twice with 2% uranyl acetate for 60 s and blotted.

100-120 micrographs were collected for each sample on Talos F200c transmission electron microscope with a Ceta 16M CMOS camera (Thermo Scientific). Nominal magnification was x45000 for 1F10 dataset (pixel size 2.30 Å), and x57000 for the other datasets (pixel size 1.8 Å). The defocus was set at -5 µm for the 1F10 dataset, and at -1.5 µm for other datasets.

Data were first processed using SIMPLE [50]. The micrographs had their contrast transfer functions estimated, and particles were automatically picked based on 2D references generated from manually picked particles. 10,000 to 30,000 particles were selected after two rounds of 2D classification and selected to build initial 3D models using SIMPLE. The particles were then exported to RELION [51, 52] followed by a round of 2D classification. Around 10k particles were selected for 3D classification using the initial 3D model generated from SIMPLE. 3D refinement was then conducted on one of the classes from the 3D classification.

#### Immunisation of mice

Animal experiments and procedures were performed according to the UK Animals (Scientific Procedures) Act Project License (PA7D20B85) and approved by the Oxford University Local Ethical Review Body. Six to eight weeks old female CD-1 mice (Envigo RMS Inc.), housed in specific-pathogen free environments, were vaccinated with full-length Pfs48/45, Pfs48/45-D1+2, or Pfs25, into each leg via the intramuscular route, using a prime-boost regime with a three week interval, with blood being collected three weeks after the final dose. Immunisations were formulated in adjuvant prior to vaccination by mixing Alhydrogel® adjuvant 2% (Invivogen; 85 μg per dose) with antigen in TBS, and incubated at RT for 1 h before injection. Sera were obtained from whole blood by leaving samples overnight at 4°C to clot, followed by 10 min centrifugation at 16,000 × g in a benchtop centrifuge at room temperature.

#### Enzyme-linked immunosorbent assays

For endpoint ELISAs, Nunc-Immuno Maxisorp 96-well plates (Thermo Scientific) coated with 2 μg/ml of antigen in PBS (Sigma) overnight at 4°C. Plates were washed with PBS-Tween (0.05%) and blocked with 10% Casein Block (Thermo Scientific). Diluted samples for end point ELISAs were added to the top row of the plate in duplicate, and used to generate a three-fold serial dilution. Plates were incubated for 2 h at room temperature and then washed as before. Goat anti-mouse whole IgG conjugated to alkaline phosphatase (Sigma) was added for 1 h at room temperature. Following a final wash, plates were developed by adding *p*-nitrophenylphosphate (Sigma) at 1 mg/mL in diethanolamine buffer (Sigma) and OD_405nm_ was read on an ELx808 microtitre plate reader (Biotek). The endpoint titre is defined as the x-axis intercept of the dilution curve at an absorbance value (±three standard deviations) greater than the OD for a serum sample from a naïve mouse. If a 1:250 dilution of a serum sample did not develop a signal above background, the IgG titre in this sample was considered to be 0. Sera of a pool of mice with a strong anti Pfs48/45-D1+2 response were included on each plate as an internal control, and all plates developed until this control reached an endpoint titre of 1,350,000 – 1,650,000. The sera of mice immunised with Pfs25 was included as a negative control.

To validate the depletion of Pfs48/45-D3 specificity from purified IgG, ELISAs were conducted as above with Pfs48/45-D3 being used as the coating antigen and IgG being serially diluted threefold down the plate from an initial concentration of 0.01mg/mL. Raw OD405_nm_ is presented. Pfs25 specific IgG was included as a negative control.

#### Standard membrane feeding assays

The ability of vaccine-induced antibodies to block the development of *P. falciparum* strain NF54 was evaluated by SMFA as previously described [53]. The percentage of mature stage V gametocytes was adjusted to 0.15 ± 0.05% and the male-female ratio was stable (almost always 1 male: 2–3 female). These were mixed with purified IgG at the concentrations (diluted in PBS) shown in the figures and then fed to 4–6 days old starved female *Anopheles stephensi* (SDA 500) via a parafilm membrane. The mosquitoes were maintained at 26°C and 80% relative humidity. After 8 days, midguts from 20 mosquitoes per group were dissected, oocysts counted, and the number of infected mosquitoes recorded. Percent reduction in infection intensity was calculated relative to the respective control IgG tested (Normal mouse IgG) in the same assay. The monoclonal antibody 4B7, specific for Pfs25 was included as a positive control.

## Statistical Analysis

Comparison of endpoint ELISA data between sera of the two groups of mice was performed by a Mann-Whitney test. The 95% confidence intervals (95% CI), and p-values of SMFA results were calculated using a zero-inflated negative binomial random effects model described previously [54].

## Data Availability

Crystallographic data and coordinates have been deposited in the PDB under the accession codes 7ZWF, 7ZWI, 7ZWM, 7ZXF and 7ZXG. Additional data supporting the findings reported in this manuscript are available from the corresponding authors on request.

## Acknowledgements

This work was funded by a Medical Research Council project grant (MR/R001138/1) to SB and MKH. This work was funded in part by the European Union’s Horizon 2020 research and innovation programme (Grant Agreement No. 733273). MKH is a Wellcome Investigator. The SMFA activity was supported by the intramural program of the National Institute of Allergy and Infectious Disease / NIH. The authors would like to thank Ed Lowe and the beamline scientists at beamlines I03, I04 and I04.1 at the Diamond synchrotron for help with crystallographic data collection. Electron microscopy data was collected at the COSMIC facility in Oxford and we thank Teige Matthews-Palmer, Rishi Matadeen and Joseph Caesar for support. We are grateful to Simon Draper and Lloyd King for facilitating use of the project license for the animal work.

## Author Contributions

K.K.,F.L., D.M., S.B and M.K.H. conceived and planned the study and wrote the manuscript. D.M. and D.D. produced Pfs48/45 and fragments. A.M. performed animal studies. D.D. and D.M. conducted endpoint ELISAs. K.M. and C.A.L. performed SMFAs. M.J. provided 32F3 and 85RF45.1. F.L. prepared Fab fragments, purified and crystallised complexes and collected X-ray diffraction data. K.K. prepared scFv fragments purified and crystallised complexes and collected X-ray diffraction data. K.K., F.L. and M.K.H. built, refined and analysed structures. F.L. performed and analysed SPR experiments. B.G. performed and analysed molecular dynamics simulations. K.K performed negative stain sample preparation, data collection and analysis. All authors read and commented on the manuscript.

## Competing interests

The authors declare no competing interests.

**Supplementary Figure 1:**
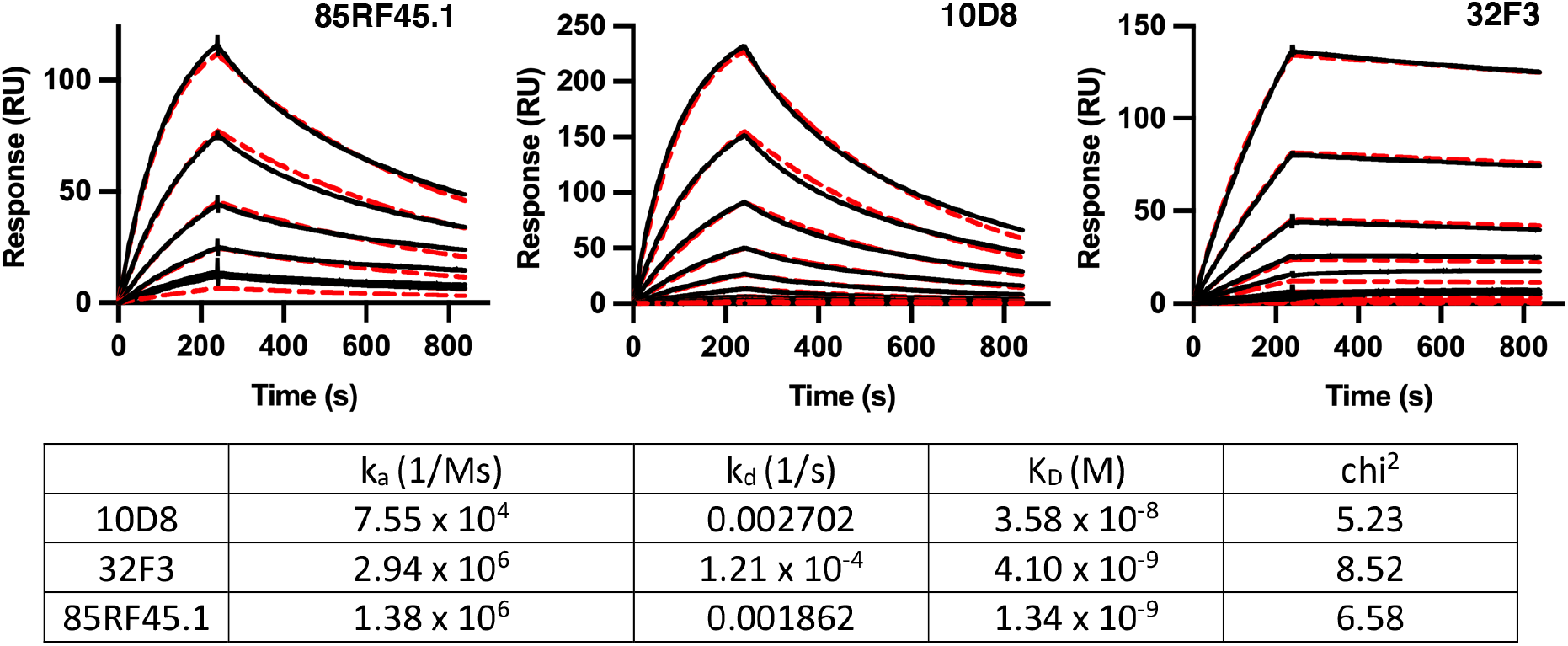
surface plasmon resonance analysis. The top panel shows surface plasmon resonance traces for binding to immobilised Pfs48/45 for antibodies 85RF45.1 (a two-fold dilution series from a top concentration of 7.8nM), 10D8 and 32F3 (each two-fold concentrations series from top concentrations of 125nM). The red dotted lines show the fitting to a one-to-one binding model. The lower panel shows the binding kinetics for these three sets of data based on the one-to-one fitting.

**Supplementary Figure 2:**
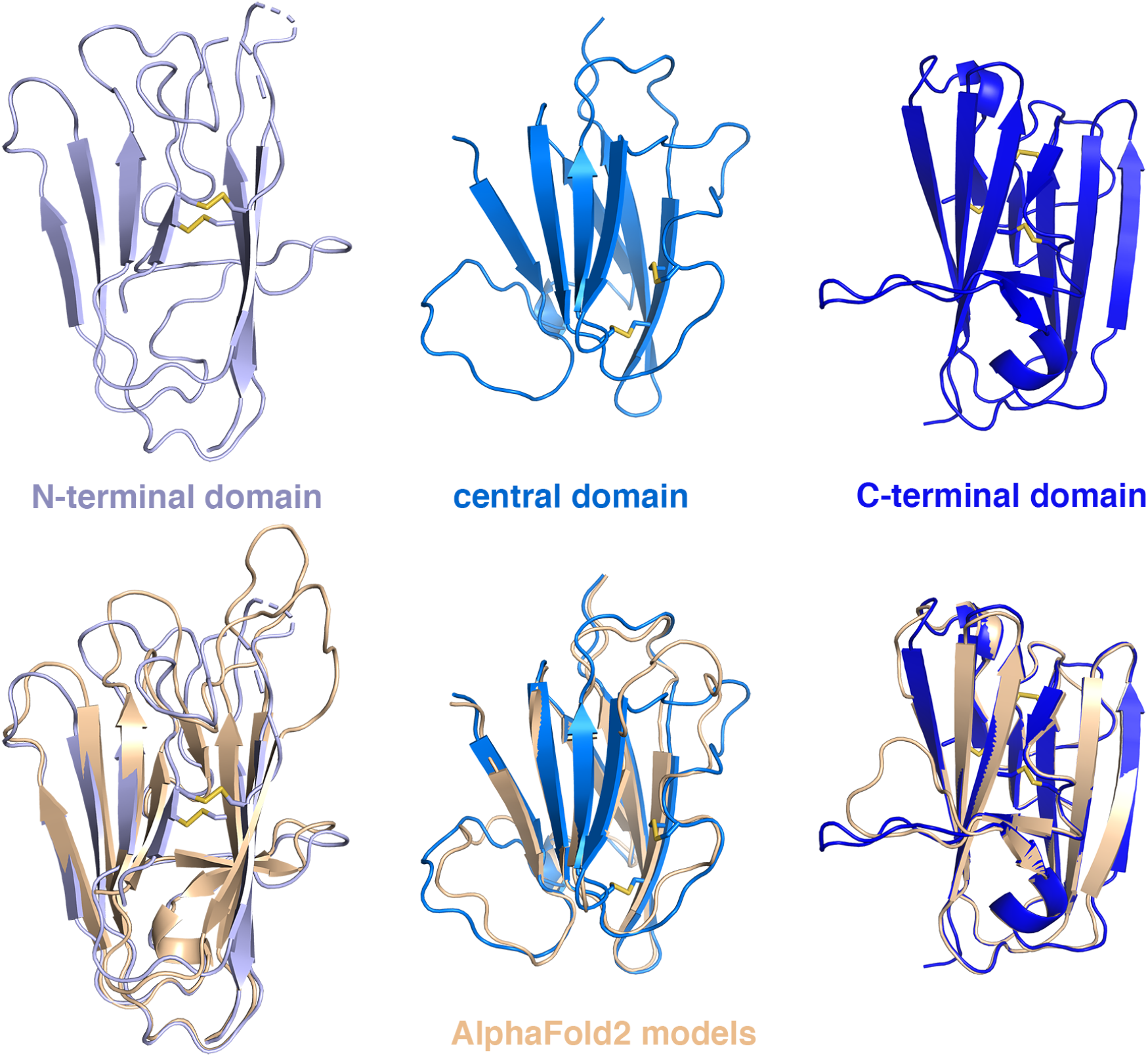
the domains of Pfs48/45 determined by x-ray crystallography compared with AlphaFold2 models. The upper panels show the N-terminal, central and C-terminal domains of Pfs48/45. Disulphide bonds are shown as sticks, with sulphur residues coloured yellow. The lower panel shows the same three domains aligned with the equivalent AlphaFold2 model in wheat colour.

**Supplementary Figure 3:**
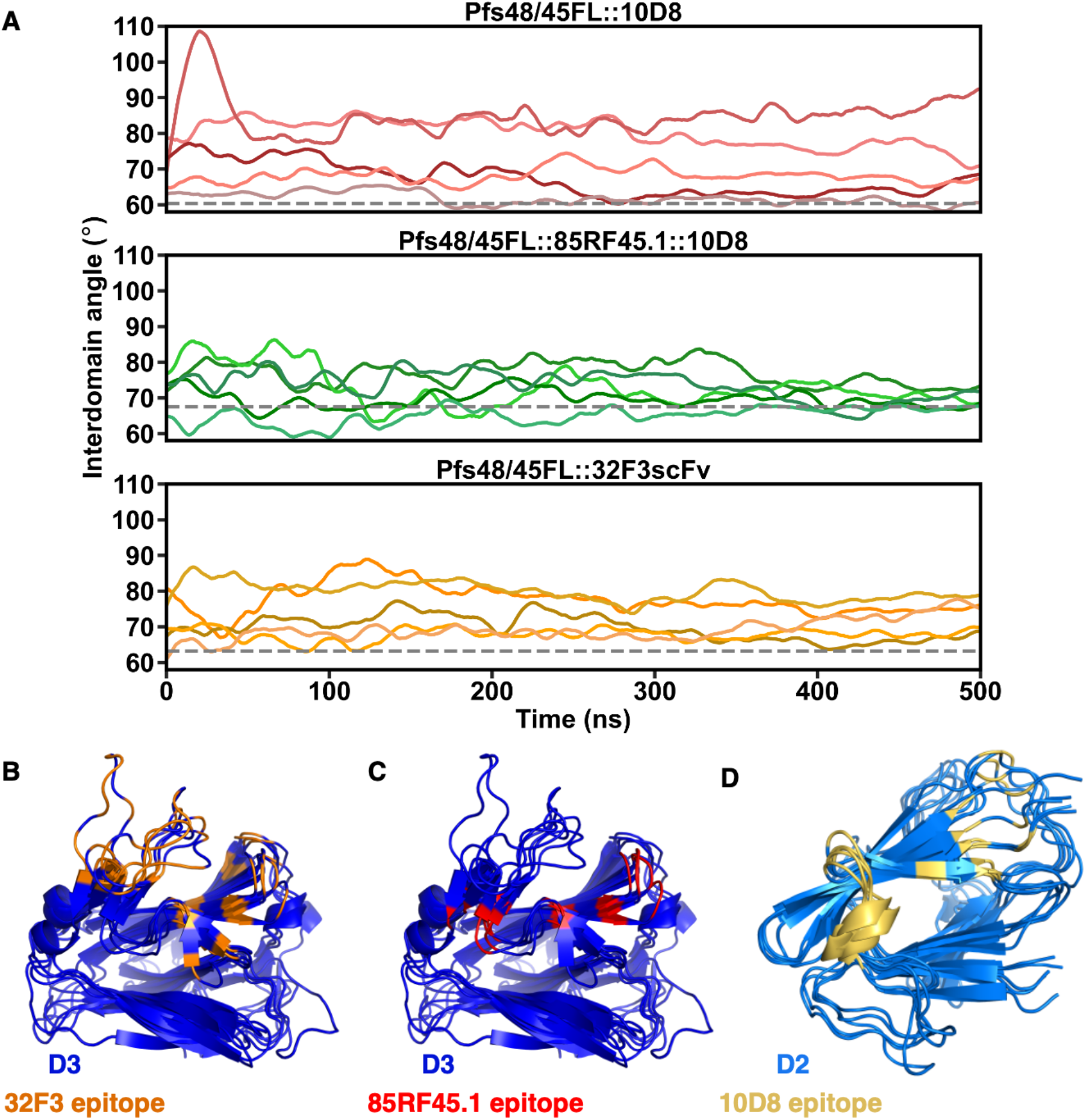
molecular dynamics simulations. **A**. Graphs showing the observed interdomain angles in full-length Pfs48/45 during atomistic molecular dynamics simulations. Three independent sets of simulations were run, using full-length Pfs48/45 taken from the crystal structure of the complex indicated above each graph. The interdomain angle is that between the line linking the centres of mass of the N-terminal and central domains and that linking the central and C-terminal domains. The five lines on each graph show five independent simulations and were smoothened using a Savistky-Golay filter. **B**. A representation of the degree of flexibility of the epitopes for 32F3 (left), 85RF45.1 (centre) and 10D8 (right). In each case, the structure of Pfs48/45 is shown in blue with the residues which contact the antibody in orange, red or yellow. Five structures are shown, from the simulations shown in **A**, to reveal the degree of epitope flexibility.

**Supplementary Figure 4:**
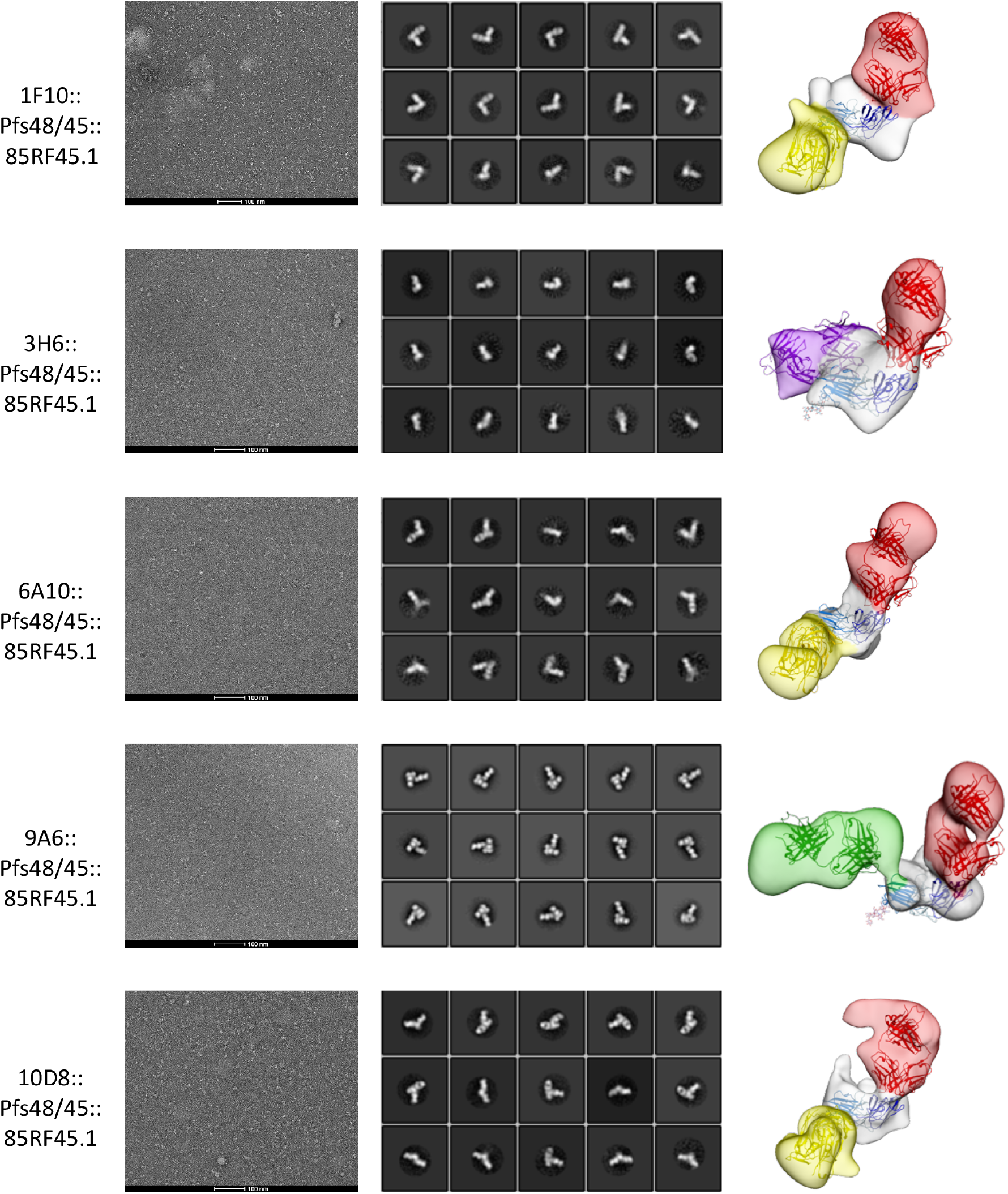
negative stain electron microscopy. Electron microscopy imaging of complexes containing Pfs48/45 (white), the Fab fragment of 85RF45.1 (red) and one additional antibody Fab fragment, as indicated to the left-hand side of each row. Each row shows a representative micrograph (left), two-dimensional class averages (centre) and a low-resolution reconstruction (right).

**Supplementary Table 1:** crystallographic statistics for structures with incomplete Pfs48/45 constructs.

| **Data collection** | **Pfs48/45_6C:32F3** | **Pfs48/45_D2+3:10D8:32F3** |
| --- | --- | --- |
| Space group | P12<sub>1</sub>1 | P2<sub>1</sub>2<sub>1</sub>2<sub>1</sub> |
| Cell dimensions: |  |  |
| a, b, c | 78.66, 78.66, 110.62 | 108.58, 158.45, 186.16 |
| $\alpha$, $\beta$, $\gamma$ | 90.0°, 99.5°, 90.0° | 90.0°, 90.0°, 90.0° |
| Resolution | 77.57 – 1.90 Å (1.93 – 1.90Å) | 108.58 – 3.69 Å (3.75 – 3.69Å) |
| Total observations | 781771 (36476) | 230227 (11072) |
| Total unique | 116014 (5704) | 35355 (1713) |
| R<sub>pim</sub> | 0.037 (0.282) | 0.065 (0.338) |
| CC<sub>1/2</sub> | 0.999 (0.780) | 0.971 (0.438) |
| I/$\sigma$(I) | 11.6 (2.2) | 7.2 (1.8) |
| Completeness (%) | 99.9 (97.8) | 100.0 (98.0) |
| Multiplicity | 6.7 (6.4) | 6.5 (6.5) |
| Wilson B factor (Å<sup>2</sup>) | 27.4 | 118.8 |
| **Refinement** |  |  |
| Number of reflections | 115991 | 35298 |
| R<sub>work</sub> / R<sub>free</sub> | 0.189 / 0.217 | 25.5 / 28.0 |
| Average B factor (Å<sup>2</sup>) | 42.0 | 161.4 |
| Number of residues: |  |  |
| Amino acid residues | 1123 | 2179 |
| Sugars | 3 | 10 |
| Waters | 963 | 0 |
| RMSZ deviations |  |  |
| Bond lengths | 0.009 | 0.004 |
| Bond angles | 1.02 | 0.67 |
| Ramachandran plot |  |  |
| Favoured | 98.5 % | 94.8 % |
| Allowed | 1.5 % | 5.2 % |
| Outliers | 0 % | 0 % |

**Supplementary Table 2:** crystallographic statistics for structures with complete Pfs48/45 constructs.

| **Data collection** | **Pfs48/45_FL:10D8** | **Pfs48/45_FL:10D8:**<br>**85RF45.1** | **Pfs48/45_FL:**<br>**32F3 scFv** |
| --- | --- | --- | --- |
| Space group | P2<sub>1</sub>22<sub>1</sub> | P4<sub>1</sub>2<sub>1</sub>2 | C222<sub>1</sub> |
| Cell dimensions: |  |  |  |
| a, b, c | 81.77, 125.80,<br>216.60 | 156.88, 156.80,<br>148.76 | 81.83, 126.12,<br>146.79 |
| $\alpha$, $\beta$, $\gamma$ | 90.0°, 90.0°, 90.0° | 90.0°, 90.0°, 90.0° | 90.0°, 90.0°, 90.0° |
| Resolution | 76.27 – 4.20 Å<br>(4.70 – 4.20Å) | 108.00 – 3.72 Å<br>(4.06 – 3.72Å) | 73.39 – 2.13 Å (2.17<br>– 2.13Å) |
| Total observations | 109594 (31747) | 456518 (56247) | 586552 (28745) |
| Total unique | 16968 (4732) | 17015 (1893) | 42648 (2277) |
| R<sub>pim</sub> | 0.088 (0.668) | 0.027 (0.875) | 0.037 (2.26) |
| CC<sub>1/2</sub> | 0.998 (0.543) | 0.999 (0.377) | 0.999 (0.384) |
| I/$\sigma$(I) | 7.5 (1.6) | 15.7 (1.2) | 11.2 (0.4) |
| Completeness (%) | 99.9 (100.0) | 94.9 (74.3) | 99.8 (99.4) |
| Multiplicity | 6.5 (6.7) | 26.8 (29.7) | 13.8 (14.2) |
| Wilson B factor (Å<sup>2</sup>) | 166.4 | 190.8 | 58.6 |
| **Refinement** |  |  |  |
| Number of reflections | 16933 | 16936 | 40794 |
| R<sub>work</sub> / R<sub>free</sub> | 25.9 / 27.6 | 28.1 / 30.2 | 27.6 / 28.6 |
| Average B factor (Å<sup>2</sup>) | 212 | 215 | 98.0 |
| Number of residues: |  |  |  |
| Amino acid residues | 1574 | 1190 | 497 |
| Sugars | 15 | 8 | 2 |
| Waters | 0 | 0 | 94 |
| RMSZ deviations |  |  |  |
| Bond lengths | 0.003 | 0.003 | 0.008 |
| Bond angles | 0.661 | 0.593 | 0.99 |
| Ramachandran plot |  |  |  |
| Favoured | 91.9 % | 92.8 % | 92.8 % |
| Allowed | 8.1 % | 7.2 % | 7.2 % |
| Outliers | 0 % | 0 % | 0 % |

**Supplementary Table 3.**
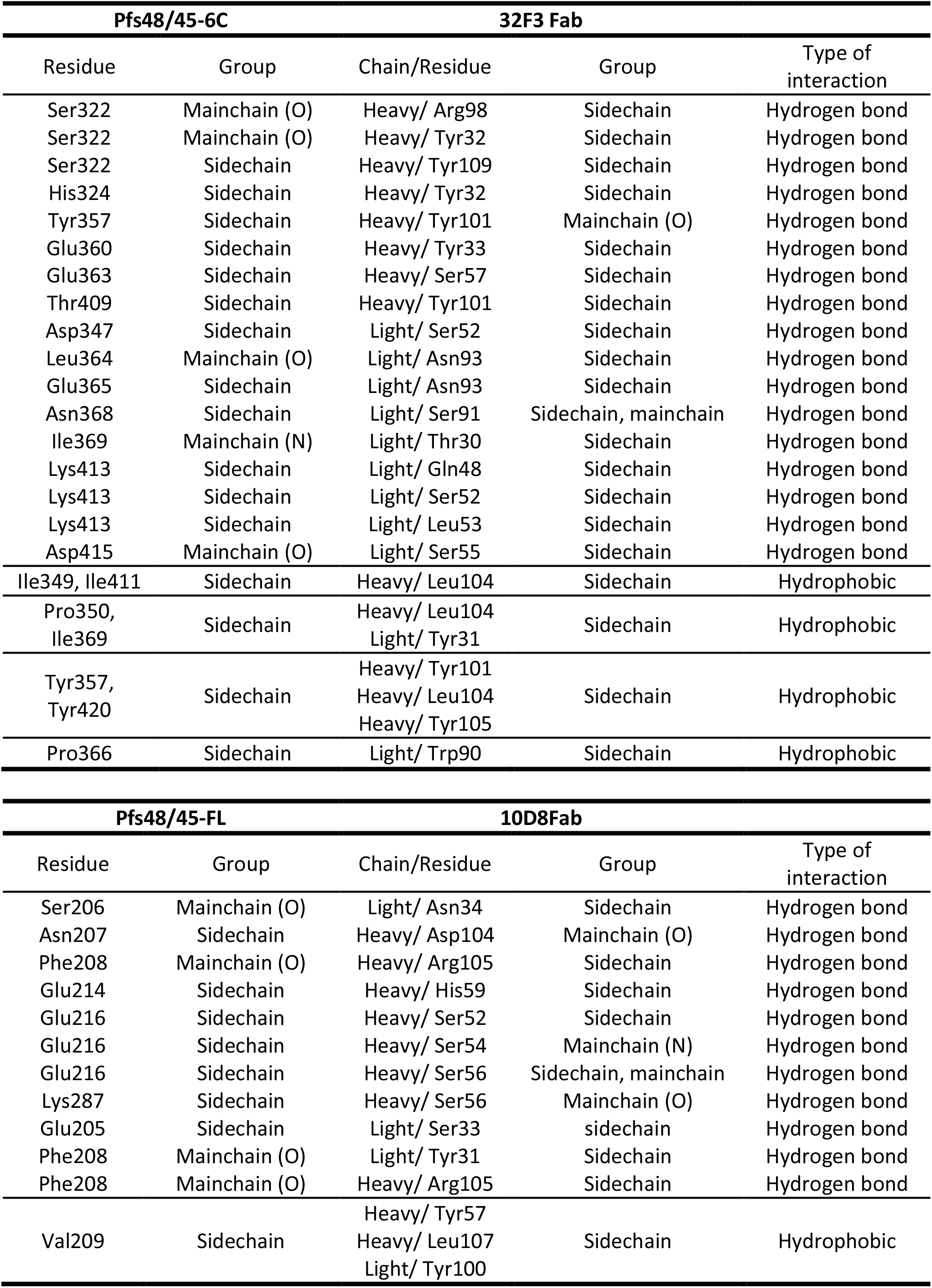
Interactions between Pfs48/45 and antibodies 32F3 and 10D8.

| Pfs48/45-6C |  |  | 32F3 Fab |  |
| --- | --- | --- | --- | --- |
| Residue | Group | Chain/Residue | Group | Type of interaction |
| Ser322 | Mainchain (O) | Heavy/ Arg98 | Sidechain | Hydrogen bond |
| Ser322 | Mainchain (O) | Heavy/ Tyr32 | Sidechain | Hydrogen bond |
| Ser322 | Sidechain | Heavy/ Tyr109 | Sidechain | Hydrogen bond |
| His324 | Sidechain | Heavy/ Tyr32 | Sidechain | Hydrogen bond |
| Tyr357 | Sidechain | Heavy/ Tyr101 | Mainchain (O) | Hydrogen bond |
| Glu360 | Sidechain | Heavy/ Tyr33 | Sidechain | Hydrogen bond |
| Glu363 | Sidechain | Heavy/ Ser57 | Sidechain | Hydrogen bond |
| Thr409 | Sidechain | Heavy/ Tyr101 | Sidechain | Hydrogen bond |
| Asp347 | Sidechain | Light/ Ser52 | Sidechain | Hydrogen bond |
| Leu364 | Mainchain (O) | Light/ Asn93 | Sidechain | Hydrogen bond |
| Glu365 | Sidechain | Light/ Asn93 | Sidechain | Hydrogen bond |
| Asn368 | Sidechain | Light/ Ser91 | Sidechain, mainchain | Hydrogen bond |
| Ile369 | Mainchain (N) | Light/ Thr30 | Sidechain | Hydrogen bond |
| Lys413 | Sidechain | Light/ Gln48 | Sidechain | Hydrogen bond |
| Lys413 | Sidechain | Light/ Ser52 | Sidechain | Hydrogen bond |
| Lys413 | Sidechain | Light/ Leu53 | Sidechain | Hydrogen bond |
| Asp415 | Mainchain (O) | Light/ Ser55 | Sidechain | Hydrogen bond |
| Ile349, Ile411 | Sidechain | Heavy/ Leu104 | Sidechain | Hydrophobic |
| Pro350, Ile369 | Sidechain | Heavy/ Leu104<br>Light/ Tyr31 | Sidechain | Hydrophobic |
| Tyr357, Tyr420 | Sidechain | Heavy/ Tyr101<br>Heavy/ Leu104<br>Heavy/ Tyr105 | Sidechain | Hydrophobic |
| Pro366 | Sidechain | Light/ Trp90 | Sidechain | Hydrophobic |
| Pfs48/45-FL |  |  | 10D8Fab |  |
| Residue | Group | Chain/Residue | Group | Type of interaction |
| Ser206 | Mainchain (O) | Light/ Asn34 | Sidechain | Hydrogen bond |
| Asn207 | Sidechain | Heavy/ Asp104 | Mainchain (O) | Hydrogen bond |
| Phe208 | Mainchain (O) | Heavy/ Arg105 | Sidechain | Hydrogen bond |
| Glu214 | Sidechain | Heavy/ His59 | Sidechain | Hydrogen bond |
| Glu216 | Sidechain | Heavy/ Ser52 | Sidechain | Hydrogen bond |
| Glu216 | Sidechain | Heavy/ Ser54 | Mainchain (N) | Hydrogen bond |
| Glu216 | Sidechain | Heavy/ Ser56 | Sidechain, mainchain | Hydrogen bond |
| Lys287 | Sidechain | Heavy/ Ser56 | Mainchain (O) | Hydrogen bond |
| Glu205 | Sidechain | Light/ Ser33 | sidechain | Hydrogen bond |
| Phe208 | Mainchain (O) | Light/ Tyr31 | Sidechain | Hydrogen bond |
| Phe208 | Mainchain (O) | Heavy/ Arg105 | Sidechain | Hydrogen bond |
| Val209 | Sidechain | Heavy/ Tyr57<br>Heavy/ Leu107<br>Light/ Tyr100 | Sidechain | Hydrophobic |

